# Dissecting contributions of directional and balancing selection to trajectories of mitochondrial haplotype evolution in *Drosophila melanogaster*

**DOI:** 10.1101/2025.05.12.653591

**Authors:** Jade R Kannangara, Tim Connallon, André N Alves, Damian K Dowling

## Abstract

Emerging evidence suggests mtDNA haplotypes contribute to fitness variation and local adaptation, with studies highlighting the potential for directional selection, imposed by distinct thermal environments, and negative frequency-dependent selection to shape spatial patterns of mtDNA haplotype diversity. However, the interplay between these different forms of selection remains unexplored. To address this, we conducted experimental evolution using *Drosophila melanogaster* populations derived from opposite ends of a latitudinal cline in Australia (Melbourne and Townsville), exposing them to contrasting thermal environments and varying starting frequencies of two mtDNA haplotypes (A1 and B1) that each segregate at high frequencies in natural populations. We paired this approach with population genetic simulations to estimate selection and its influence on haplotype evolutionary trajectories. We found that mtDNA haplotype frequencies were influenced by complex interactions among temperature, starting frequency, and nuclear genomic background (Melbourne, Townsville, or admixed). Simulations supported directional selection as a driver of haplotype frequencies in populations evolving at 17°C in the Melbourne background, and otherwise selection was predominantly balancing and shaped by negative frequency-dependent fitness effects of the haplotypes. Finally, our results suggest that the thermal environment is not the sole environmental variable shaping mitochondrial evolutionary dynamics and point to a possible role for mito-nuclear epistasis.

## Introduction

In eukaryotes, mitochondria are crucial for cellular respiration, a process that requires precise coordination between gene products encoded by the mitochondrial and nuclear genomes (Rand *et al*. 2004; Wolff *et al*. 2014). The mitochondrial genome (mtDNA) encodes a small number of genes involved in this process. These genes are highly conserved, particularly among animals, which (with rare exceptions) carry the same core set of 37 mtDNA genes. Given the small number and the critical and conserved functions of mitochondrial genes, it was long assumed that functionally important mtDNA sites would be subject to strong purifying selection and the mitochondrial genome would therefore contribute very little to the genetic variation of fitness components or adaptive evolutionary divergence over time and between species. Contrary to this assumption, numerous studies over the last two decades have revealed that mitochondrial sequence variation has surprisingly large effects on various traits associated with fitness, including metabolism, physiology, morphology and life-history (Rand 2001; Dowling *et al*. 2008; Burton *et al*. 2013; Ballard and Pichaud 2014; Dobler *et al*. 2014; Wallace 2016; Hill *et al*. 2019; Dowling and Wolff 2023). Population genetic analyses of the mitochondrial genome also show that a substantial fraction of mtDNA substitutions are adaptive and fixed in response to natural selection (Bazin *et al*. 2006; James *et al*. 2016).

Several recent studies have reported geographic clines in mitochondrial haplotype frequencies associated with latitude, longitude or altitude (Grant *et al*. 2006; Cheviron and Brumfield 2009; Silva *et al*. 2014; Consuegra *et al*. 2015; Camus *et al*. 2017; Wada *et al*. 2019; Kapun *et al*. 2020), raising the intriguing possibility that the mitochondrial genome might disproportionately (given its size) contribute to local adaptation. Examples from vertebrates include geographic associations between mtDNA haplotypes or patterns of mtDNA variation and major climate zones in both humans and birds (Lamb *et al*. 2018), where different haplotypes might be favoured (Mishmar *et al*. 2003; Ruiz-Pesini *et al*. 2004; Balloux *et al*. 2009); evidence for adaptive divergence of mtDNA protein sequence variants across latitudinal gradients of Atlantic salmon and European anchovies (Silva *et al*. 2014; Consuegra *et al*. 2015), and mtDNA haplotype frequency clines across elevational gradients of the rufous-collard sparrow (Cheviron and Brumfield 2009). A well-studied example from an invertebrate (and our focus here) involves a pair of mtDNA haplotypes of the fruit fly *Drosophila melanogaster*. Haplotypes A1 and B1, which differ by 15 single nucleotide polymorphisms, exist on a latitudinal cline along the east coast of Australia, where A1 haplotype frequencies have been observed to increase as the latitude decreases (Camus *et al*. 2017).

While patterns of clinal divergence suggest that mitochondrial genomes might contribute to local adaptation, allele frequency clines can also arise from nonadaptive processes associated with historical demographic events (Adrion *et al*. 2015). Moreover, even in cases where demographic explanations can be excluded, identifying the environmental or ecological variables that underlie selection for clinal divergence remains a challenge, as many biotic and climatic variables (co)vary across species’ ranges, only some of which influence selection (Wade and Kalisz 1990; Kawecki and Ebert 2004). Controlled experiments are therefore essential for evaluating whether clinal divergence is adaptive and identifying the environmental variables that influence selection in nature (e.g., (Caruso *et al*. 2017; Carvalho *et al*. 2022)). Controlled lab studies have been particularly useful for testing whether the spatial distributions of mtDNA haplotype frequencies are consistent with local adaptation to thermal conditions (Camus *et al*. 2017; Lajbner *et al*. 2018). Immonen *et al*. (2020) used experimental evolution of seed beetle populations (*Callosobruchus maculatus*) exposed to different temperatures and observed changes in mtDNA haplotypes frequencies that were consistent with thermal adaptation (Immonen *et al*. 2020). Camus *et al*. (2017) established *D. melanogaster* strains carrying the A1 or B1 mtDNA haplotypes, alongside a standardized nuclear genetic background, and tested whether these haplotypes affect thermal stress resistance. Consistent with the natural spatial distribution of these haplotypes, the A1-carrying strains exhibited greater resistance to heat and lower resistance to cold stresses than B1 strains. However, these findings were not evident when the mtDNA haplotypes were tested in genetically variable ‘mass-bred’ nuclear backgrounds (Bettinazzi *et al*. 2024). Notwithstanding this, Lajbner *et al*. (2018) used an experimental evolution approach in mass-bred populations initiated with equal starting frequencies of the two haplotypes and found that A1 haplotypes increased in frequency in warm environments (averaging 25°C) while B1 increased in cooler environments (averaging 19°C). Intriguingly, these temperature-dependent changes in haplotype frequency were only observed in populations that had been cured of *Wolbachia* infections via antibiotic treatment (Lajbner *et al*. 2018).

Combined, these studies generally support the hypothesis that selection favours different mitochondrial haplotypes in different thermal environments (e.g., A1 haplotypes favoured in warm and B1 in cold environments in *D. melanogaster*). Nevertheless, the specific forms of selection affecting mtDNA variation in different environments is unclear. Most studies appear to (at least tacitly) assume that selection in each environment is directional and, in the absence of gene flow, would result in fixation of a single locally favoured mtDNA haplotype. An alternative is that selection in some thermal environments might maintain multiple genetic variants rather than favouring the fixation of any single one (i.e., “balancing selection” might maintain mtDNA diversity). Multiple distinct mitochondrial haplotypes are often observed to segregate at intermediate frequencies within wild populations (González *et al*. 1994; Hansen and Loeschcke 1996; García-Martínez *et al*. 1998; Oliver *et al*. 2005; Camus *et al*. 2017; Ueno and Takahashi 2021), which could be explained by balancing selection, though other hypotheses—including selective neutrality, or high gene flow among populations favouring different mtDNA haplotypes—might also account for such patterns.

There are various forms of balancing selection, but most apply to standard Mendelian systems of inheritance. The uniparental inherence of mitochondrial genes renders balancing selection unlikely unless the relative fitness associated with each mtDNA haplotype is negative frequency-dependent (NFD), with rare variants selectively favoured over common ones (Gregorius and Ross 1984; Kazancıoğlu and Arnqvist 2014; Arnqvist *et al*. 2016). Consistent with this possibility, Oliver *et al*. (2005) observed that two mtDNA haplotypes were maintained at intermediate frequencies for over 30 generations in replicated laboratory populations of *D. subobscura* (Oliver *et al*. 2005), which a subsequent simulation study suggested is consistent with a NFD model of balancing selection (Arnqvist *et al*. 2016). In another experimental evolution study using seed beetles, Kazancioǧlu and Arnqvist (2014) showed that rare mtDNA haplotypes were consistently favoured over common ones, implying that NFD fitness effects lead to balancing selection of the haplotypes.

These studies highlight the potential for frequency-dependence, directional selection, and/or balancing selection, to influence mitochondrial haplotype diversity in different thermal environments. However, the interplay of these factors remains unclear. To address this gap, we carried out experimental evolution of *D. melanogaster* laboratory populations to estimate selection and its influence on mitochondrial haplotype evolution in different thermal environments. We specifically investigated whether the outcomes of thermal selection were contingent on the nuclear genomic background and the initial population frequencies of mtDNA haplotypes. We studied populations harbouring the A1 and B1 mtDNA haplotypes previously studied by Camus et al (2017) and Lajbner *et al*. (2018) and varied their initial proportions to test for frequency-dependent evolutionary responses in cold versus warm environments. Each population’s nuclear genomic background was either entirely derived from flies collected from a cline-end population (Melbourne or Townsville) or it was heterogeneous due to admixing flies with Melbourne and Townsville nuclear backgrounds. The mtDNA frequencies were tracked across 10 generations of experimental evolution, with effects of thermal environment, initial haplotype frequency, nuclear genomic background, and their interactions estimated in a generalised linear mixed model. Finally, we used population genetic simulations mirroring our experimental design and Approximate Bayesian Computation to identify selection parameters that were consistent with our experimental data.

## Materials and Methods

### Creation of mito-nuclear strains and fly husbandry

Flies harbouring the A1 and B1 haplotypes were collected from Coffs Harbour in 2016, which is located at the approximate midpoint of the well-studied latitudinal distribution of *Drosophila* in eastern Australia (Hoffmann *et al*. 2001; Arthur *et al*. 2008; Telonis-Scott *et al*. 2011; Camus *et al*. 2017; Lajbner *et al*. 2018). Individual wild-mated females became founders of separate isofemale lines (Ekta Kochar, Damian Dowling, unpublished data). Following custom mitochondrial genotyping, two of these lines were chosen to be the mitochondrial donors for a set of *mito-nuclear strains*, which are described below. In July 2021, each isofemale line was treated for three generations with tetracycline hydrochloride (concentration 0.3mg/ml of potato-dextrose medium) to ensure effective removal of *Wolbachia*.

The lab populations contributing nuclear backgrounds to the mito-nuclear strains were collected from Melbourne [-37.77,144.99] and Townsville [-19.26, 146.81] between December 2020 and March 2021. These populations were chosen as they are located near cline-ends in which the A1 and B1 haplotypes naturally segregate. Twenty-six isofemale lines collected from Melbourne were used to create a Melbourne mass-bred population (12 virgin females and 12 virgin males from each line were admixed to form one large panmictic population) and thirty-five isofemale lines collected from Townsville were used to create the Townsville mass-bred population (10 virgin females and 10 virgin males from each line were admixed). Two replicates of each population were created in this manner. These outbred populations were treated with tetracycline for two generations to remove *Wolbachia* (concentration 0.3mg/ml of potato-dextrose medium) and each population was maintained across two 300mL bottles carrying 400 flies each. For each replicate, the two bottles were mixed each generation and then redistributed back out into 2 bottles, where 5-day post-eclosion females were allowed to lay eggs for 24 hours to propagate the next generation.

Beginning August 2021, A1 or B1 haplotypes were substituted into each of the mass-bred population backgrounds to create four combinations of mtDNA × nuclear genetic background. Crosses were carried out for six generations using the balancer-chromosome scheme described in Clancy (2008) (Figure S1), resulting in females carrying a donor mitochondrial haplotype (A1 or B1) and a nuclear genetic background from one of the mass-bred populations (the four combinations were created in replicate to form eight mito-nuclear strains). Each mito-nuclear strain is severely bottlenecked during its creation, due to small numbers of flies possessing the required genotypes during the balancer chromosome crossing scheme. Following completion of the scheme, we restored representative levels of nuclear genomic variation, by backcrossing 32 virgin females from each of the created strains to 32 males of the corresponding mass-bred population for approximately ten more generations. All strains were maintained on a discrete 12hr day/night cycle at 25 degrees in 300 ml bottles containing standard potato-dextrose food medium, with live yeast added *ad libitum* to promote oviposition.

### Experimental Evolution Design

The 8 mito-nuclear strains were combined in various permutations to establish 64 *experimental populations*, which differed in the starting frequencies of A1 and B1 haplotypes, the constitution of the nuclear genomic background, and the rearing temperature. Experimental populations each founded by 500 adult flies at 1-5 days post-eclosion, comprised of an equal mix of ages and equal sex ratio. We used two different starting frequencies for the mtDNA haplotypes (30% and 70% for each), which allow for tests of frequency-dependent fitness effects of the haplotypes (Table S1). Four types of initial nuclear genomic background were used across the 64 experimental populations: (1) 100% derived from a mass-bred Melbourne population, (2) 100% from a mass-bred Townsville population, (3) admixed with 30% from Townsville and 70% from Melbourne, and (4) admixed with 70/30 from Townsville/Melbourne (see Table S1). Finally, populations were allocated to one of two contrasting temperatures (17°C or 27°C, Figure S2). Each of the 8 mito-nuclear strains was used twice to create the required combinations of experimental population (8 mito-nuclear strains × 2 temperatures × 2 technical duplicates × 2 starting haplotype frequencies = 64 populations).

Within each experimental population, adult flies cohabited for either 2 days (for 27°C treatment) or 4 days (17°C treatment) in 500mL bottles containing a standard potato-dextrose medium. Flies subjected to the 17°C treatment were allowed longer cohabitation because cooler temperatures lead to slower development and maturation of oogenic machinery. Following cohabitation, flies were transferred to 500mL bottles containing apple juice agar, supplemented with live yeast paste for either 24 h (27°C treatment) or 48 h (17°C treatment), and allowed to oviposit. Adults were then removed and frozen at -20°C for subsequent mitochondrial genotyping. Eggs were collected from the apple juice agar bottles and washed in Phosphate-Buffered Saline (PBS). Approximately 500 eggs (±35) were transferred to new 500mL bottles containing standard potato-dextrose medium, as described in Nouhaud *et al*. (2018), and allowed to develop into adults at their assigned temperatures. A pilot study showed that a 4-day oviposition period in the 17°C treatments was required for flies to generate enough eggs for the next generation, since oviposition rates are slower at low temperatures. Rearing temperature does not, however, affect the egg density per mL of PBS, as long as flies are given sufficient time to lay the required number of eggs. Following eclosion, the new generation of flies was transferred to new 500mL bottles, where they cohabited for either 2 days (27°C treatment) or 4 days (17°C treatment) to allow mating and ovipositing. This process was repeated for 10 generations, with populations maintained at their assigned temperatures under a 12hr light/dark cycle (Figure S2). In generation 2, a fungal outbreak confined to one experimental population (30% A1 Melbourne × 70% B1 Townsville – 17°C treatment) resulted in its extinction; the other 63 populations survived 10 generations of experimental evolution.

### DNA extractions and genotyping

DNA was extracted from 30 single female flies sampled from each experimental population at generations 3, 5 and 10 (30 flies × 63 populations × 3 generations = 5,670 single DNA extractions).

Each DNA extraction was based on a single female fly placed in a well of a 96-well plate. Flies were homogenised in 100uL of buffer (100mM Tris-HCl, pH 8.8, 0.5mM EDTA and 1% SDS) and plates were incubated at 65°C before 1:10 3M sodium acetate was added. Plates were then incubated on ice for 30 minutes to precipitate the protein and SDS before being centrifuged at full speed for 15 mins at 4°C. Afterwards, 40uL of supernatant was removed from each well and transferred to a new well on a 96-well plate, and 20uL of isopropanol was added to each well to precipitate DNA. After centrifugation at full speed for 5mins at room temperature, the supernatant was discarded, and 70% ethanol was added to wash the pellet. Ethanol was then removed and 50uL of milli-Q H_2_O was added to re-suspend the pellet. The DNA was stored at -20°C until needed.

The presence of either the A1 or B1 haplotype was determined via multiplex PCR using two forward primers. The first primer, specific to the A1 haplotype (mtA1-F, with sequence: 5’TTCATTCTTGAACAGTACCTGtT 3’), was designed as per a previously described ARMS-qPCR, which is a method used to differentiate between mitochondrial haplotypes differing by a SNP (Wang *et al*., 2011). As such, a deliberate mismatch (shown in lower-case) was included in the second to last base on the 3’ end of the primer to increase binding specificity in our multiplex PCR. The second forward primer (with sequence 5’CAGGTTTTATTCACTGATACC 3’) was common between the A1 and B1 haplotypes (mtcontrol-F) and served as a control. We used one reverse primer (sequence 5’ TCGTCCAGGTGTACCGTCAAC 3’) that was common to both mtDNA haplotypes (mtcontrol-R). In the presence of the A1 haplotype, two bands were expected, including one A1-specific 56bp band and one control 978bp band. For the B1 haplotype, one control band of 978bp was expected.

PCR was performed using the Go-Taq Green Mastermix (Promega, M7123). Individual PCRs were performed in a 96-well plate in 12.5uL reactions where the mtA1-F, mtcontrol-F and mtcontrol-R primers were added in a 0.5:0.5:1 ratio, respectively. All primers were at a 10uM concentration. PCRs were performed in a PCR thermocycler (Eppendorf) under the following conditions: 95°C for 2 min for 1 cycle, followed by 95°C for 30secs, 50°C for 30sec and 72°C for 11 secs for 30 cycles, concluding with 72°C for 5 min for 1 cycle.

Following amplification via PCR, approximately 8uL of PCR product was run on a 2% agarose gel consisting of Red-Safe (Scientifix) at 110 volts for 30 minutes. Gels were visualised on a ChemiDoc MP Imaging system (Bio-Rad). Two positive controls consisting of known A1 and B1 DNA were also run on each gel, as well as a negative control. Genotypes for each population were scored blind to the identity of the experimental population. A maximum of 30 flies and minimum of 21 flies were scored for each sampling timepoint and experimental population, accounting for unsuccessful DNA extractions.

### Statistical Analyses

We fitted a generalised linear mixed model to the genotype data, modelling factors that affected the frequency of A1 within each population, using the *glmmTMB* package 1.1.10 (Brooks *et al*. 2017) in R 4.0.3 (R Core Team 2023). A binomial vector was created as the response variable (number of A1 and number of B1 haplotype-bearing flies per population). The fixed factors included temperature (either 17°C or 27°C), starting proportion of A1 (0.3 or 0.7), and nuclear genomic background (either Melbourne, Townsville, 70% Townsville or 30% Townsville). Generation (3, 5, 10) was modelled as a fixed variate, and interactions between these fixed effects were also included in the model. To account for hierarchical structure and overdispersion, we included random intercepts and slopes of Generation nested within population replicate (63 levels), with uncorrelated random effects, using the || specification in glmmTMB; this accounted for the fact that we had three repeated measurements (3, 5, 10) of each population (Schielzeth and Forstmeier 2009). An additional random intercept for each observation was included to model residual overdispersion (Harrison 2015). Models were fit using maximum likelihood estimation. Type III Wald chi-square tests for fixed effects were performed using the Anova() function from the *car* package (Fox and Weisberg 2019). Sum-to-zero contrasts were used for all categorical variables to ensure estimable main effects in the presence of interactions. Statistical significance was assessed at α = 0.05.

We planned to reduce the fitted model by simplifying a full model, sequentially removing terms that did not change (at α = 0.05) the deviance of the model (Fox, 2002). However, the full model was the final model, since the 4-way interaction between the fixed effects was statistically significant.

### Population genetic model and approximate Bayesian computation (ABC) analysis

We used approximate Bayesian computation (ABC) to test the fit between the data from the experimental evolution study and a simple population genetic model that includes a pair of mtDNA haplotypes and potential for frequency-dependence, directional selection, and balancing selection. The model defines fitness of A1 individuals relative to B1 individuals as 𝑤_𝐴_ = 𝑉(1 + 𝑠𝑝), where *V* denotes the relative fitness of A1 individuals in a population predominantly comprised of B1 individuals, *s* represents the strength and direction of frequency-dependence of fitness, *p* represents the frequency of the A1 haplotype, and *q* = 1 – *p* is the frequency of the B1 haplotype. The parameters *V* and *s* are subject to the constraints: 𝑉 ≥ 0 and 𝑠 ≥ −1, which ensure that fitness values must be non-negative. Values of *s* can fall into three possible domains of frequency-dependence: positive frequency-dependence (PFD, where 𝑠 > 0), negative frequency-dependence (NFD, where 0 > 𝑠 ≥ −1), and no frequency-dependence (𝑠 = 0). See Fig. 1 for biological intuition on the model parameters.

**Figure 1.**
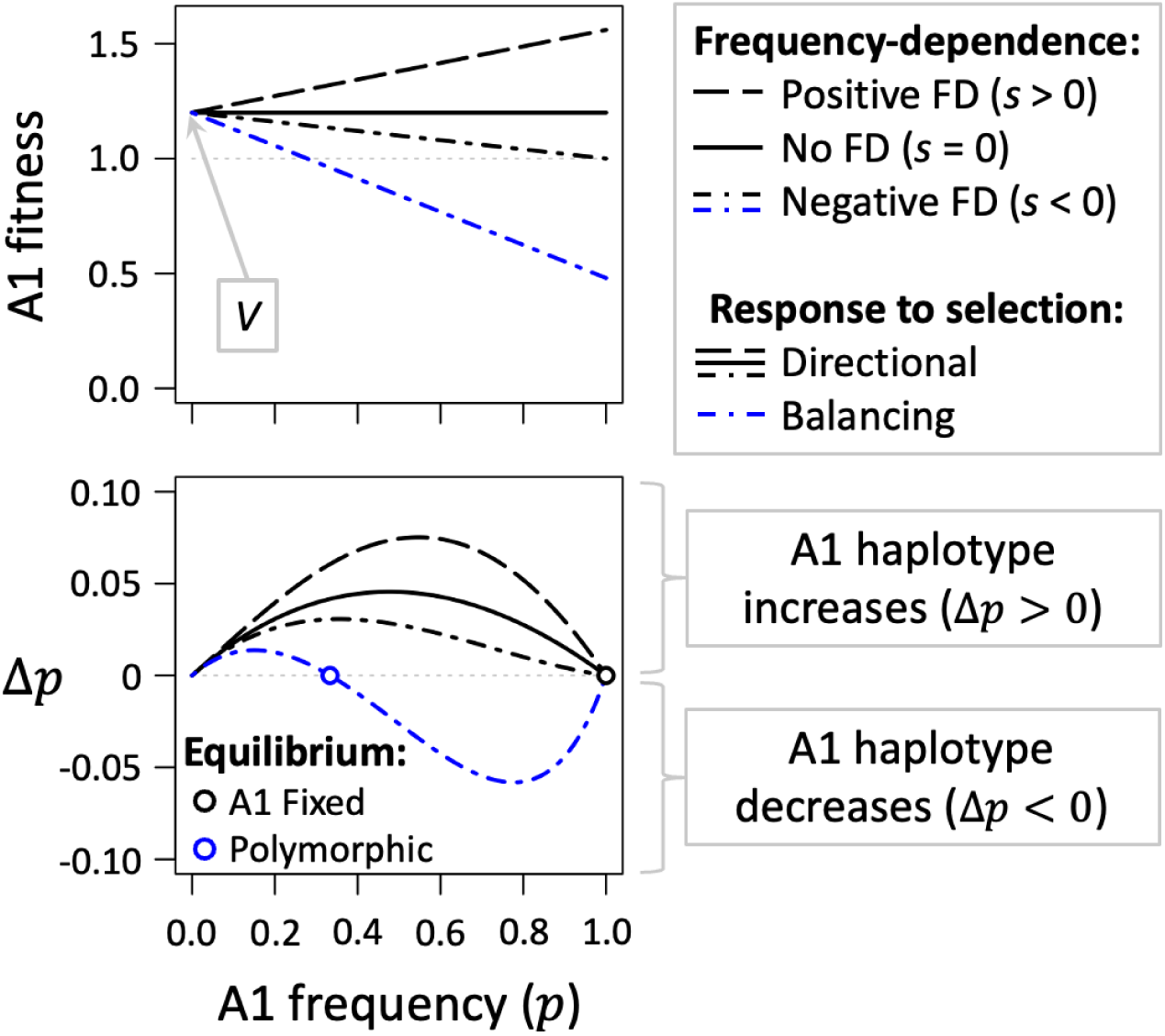
A simple model allowing for frequency-dependence of fitness, directional selection, and balancing selection of mtDNA haplotypes. In each example shown, the A1 haplotype has an advantage when rare (*i.e.*, *V* > 1) and consistently increases in frequency when rare (*i.e.*, Δ𝑝 > 0 when the A1 frequency, *p*, is small). Frequency-dependence (FD) can be positive (*s* > 0), negative (*s* < 0) or absent (*s* = 0). Sufficiently strong negative FD can lead to balancing selection (blue curves). In the remaining cases shown (in black), directional selection favours the fixation of A1. Results are shown for models with *V* = 1.2, *s* = (0.3, 0, -0.167, -0.6), and values for Δ𝑝 are based on eq. (1).

The evolutionary dynamics of the model are described by the difference equation:

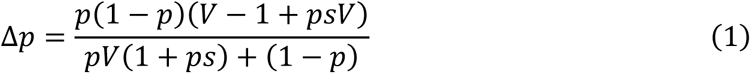

which describes the *expected change* in the A1 haplotype frequency, per generation. In the absence of drift (which we return to further below), the model has four possible long-term stable equilibrium states towards which the population is expected to evolve. When 𝑉 < 1 and 𝑉(1 + 𝑠) < 1, then A1 is always less fit than B1 and selection favours fixation of the latter (*i.e.*, there is “directional selection” for B1 and against A1). When 𝑉 > 1 and 𝑉(1 + 𝑠) > 1, then A1 is always more fit than B1, leading to directional selection in favour of A1 (see the black curves in Fig. 1). When 𝑉 > 1 and 𝑉(1 + 𝑠) < 1, then each haplotype is favoured when it is rare and “balancing selection” maintains both haplotypes (see the blue curves in Fig. 1). Finally, the system is “bi-stable” when 𝑉 < 1 and 𝑉(1 + 𝑠) > 1 (whether A1 or B1 is fixed depends on their initial frequencies).

To identify parameter combinations of the model that are consistent with our experimental results, we carried out stochastic simulations over a broad range of plausible parameter values of haplotype fitness (*V* and *s*) and effective population size, coupled with random sampling of individuals for genotyping and haplotype frequency estimation, which mimics our experimental design. Simulated haplotype frequency estimates for a given simulation were then compared to our experimental data. As in our experiments, 250 females initially founded each population. For each simulation run, we drew an effective population size for females (*N_f_*, which is relevant to mitochondrial genome evolutionary dynamics) from a uniform prior distribution between 100 and 200 (i.e., *N_f_* ∼U(100, 200)). This interval spans a plausible range for our experiments, in which the census number of breeding females per generation was 250. Selection coefficients were drawn from uniform priors: *s* ∼U(−1, 1) and *V* ∼U(0, 3). For each parameter set, we carried out Wright-Fisher forward simulations of replicate experimental evolution populations for 10 generations (four replicates per environmental treatment, excepting populations evolved at 17°C with an initial A1 frequency of 30% and an admixed 30% Townsville nuclear background, which had three replicates). For each generation, deterministic model predictions were used to calculate the expected genotype frequencies in breeding females of that generation; multinomial sampling was used to determine the actual genotype frequencies in the *N_f_* breeding females (see pp. 229-230 of Charlesworth and Charlesworth 2010).

In each simulated population, we mimicked the process of sampling and genotyping of our experiments (30 individuals each in generations 3, 5, and 10). Simulated frequency estimates per population were used to calculate the mean frequency across replicates of a given environmental treatment, per sampled generation. These estimates were compiled in a vector with six elements: three representing mean A1 frequency estimates at generations 3, 5, and 10 from populations initiated with a 30% A1 starting frequency, and three from populations initiated at 70% A1 starting frequency. We calculated Euclidean distance (*d*) between each simulated frequency vector and the corresponding vector of frequency estimates from the experiment. A simulation was rejected when the distance metric was large (*d*^2^ > 0.01) and accepted when the distance measure was small (*d*^2^ < 0.01). We recorded parameters for each accepted simulation, which were used to calculate posterior parameter distributions that were consistent with experimental data. Simulations for each model and environmental treatment were carried out until 1000 parameter sets were accepted.

## Results

Population frequency changes of the mtDNA haplotypes were influenced by an interaction between temperature, nuclear genomic background, the initial haplotype frequencies, and generation (χ^2^=12.76, p<0.01, Table 1, Figure 2). Several lower-order interactions and main effects involving these factors also contributed to the mtDNA haplotype frequencies (Table 1). During the ten generations of the experiment, the A1 haplotype increased in frequency across most of the experimental populations. However, the magnitude of increase across generations was typically larger in populations subjected to the 17°C treatment compared to those at 27°C (χ^2^=13.97, p<0.001, Table 1), and in populations with a starting A1 frequency of 0.3 rather than 0.7 (χ^2^=32.50, p<0.001, Table 1, Figure 2). When the starting frequency of A1 was 0.3, the A1 haplotype strongly increased by generation 3, with further increases across subsequent generations in most experimental populations. An exception was observed in populations with admixed nuclear genomic backgrounds of 70% Townsville and 30% Melbourne that evolved at 17°C (Figure 2). Conversely, evolutionary changes in A1 frequency were more variable in populations with a starting frequency of 0.7. At 17°C, A1 increased, except in admixed populations with 30% Townsville nuclear backgrounds, where A1 decreased to roughly *p* ∼ 0.5 and remained stable thereafter (Figure 2). At 27°C, haplotype frequencies remained relatively stable across generations.

**Figure 2:**
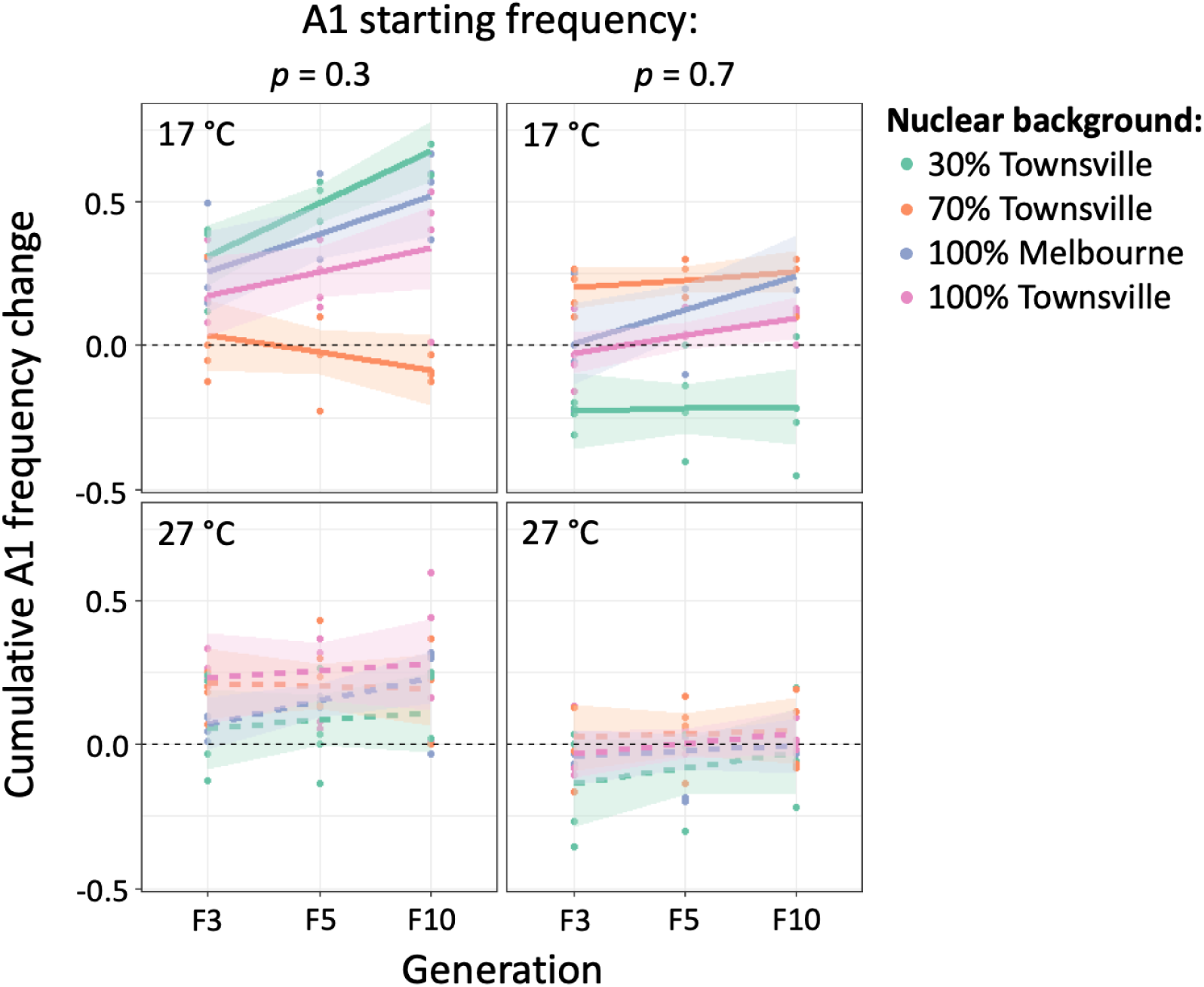
Effects of temperature, initial haplotype frequencies, and nuclear genomic background on A1 haplotype frequency change. Results show the cumulative change in A1 frequency estimated during experimental evolution (estimates were based on genotyping of samples from generations 3, 5, and 10). Dots show estimates from individual experimental populations, while regression lines (with 95% confidence intervals) show the best fit linear model for A1 frequency change in response to effects of (and interactions between) generation, starting frequency of the A1 haplotype, and temperature across four different nuclear backgrounds.

**Table 1:**
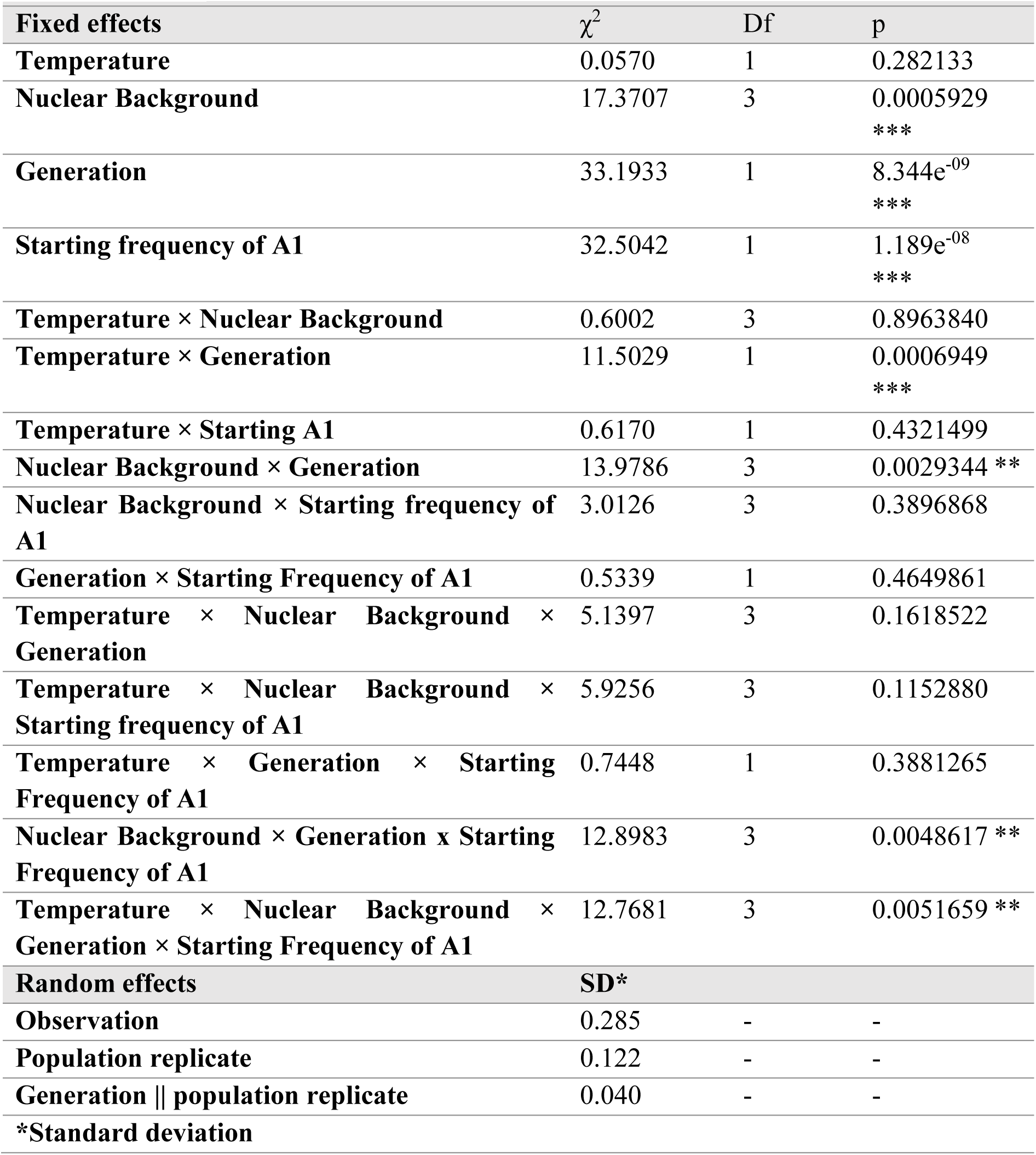
Generalised linear mixed model of sources of variance affecting A1 haplotype change. The model explores the effects of temperature, nuclear background, generation, and initial mtDNA haplotype frequency on changes of A1 haplotype.

In a few cases, a sample of genotyped individuals was fixed for one of the two mtDNA haplotypes. Fixation of A1 was observed in seven of the 63 experimental populations (∼10%), while B1 was never observed to fix. In all cases, fixation events occurred in populations evolved at 17°C. In four of the six cases, the starting frequency of A1 was 0.3, and in four of six cases, the nuclear genomic background was admixed. In at least some of these cases, A1 was only fixed in the sample and not in the population (e.g., in one case, a 5^th^-generation sample implied A1 fixation, but a later sample did not), which must be kept in mind when interpreting these observations. However, it is clear that A1 showed dramatic frequency increases in several experimental populations, particularly when temperatures were low.

Overall, these results suggest that mtDNA haplotype frequency changes were shaped by natural selection, with evolutionary outcomes varying by environment and the initial population genetic background. We further probed this through population genetic simulations, which follow.

### Approximate Bayesian estimation of selection of mtDNA haplotypes

We used Approximate Bayesian Computation (ABC) to estimate population parameters of selection and drift that are consistent with our experimental evolution data for each thermal environment and nuclear genomic background. The ABC analysis is based on a simple and flexible population genetic model (Fig. 1; described in the methods), in which the fitnesses associated with different mtDNA haplotypes can be frequency-dependent (we also allow them to be frequency-*in*dependent) and evolutionary dynamics can result in either the elimination or preservation of genetic variation. Computer simulations involving selection, genetic drift, and random sampling of individuals for genotyping, were used to generate theoretical datasets that were compared to our actual data. Rejection-sampling in the ABC framework (based in similarity criteria, as described above) was used to generate posterior distributions of three parameters: (1) the female effective population size per treatment (*N_f_*), (2) the fitness of the A1 relative to the B1 haplotype when A1 is rare (*V*; see Fig. 1), and (3) the strength and direction of frequency-dependence of mtDNA haplotype fitnesses (*s*; see Fig. 1). Since plausible values of *N_f_* did not meaningfully influence our estimates of selection, we focus below on the selection estimates. In addition, since our model was unable to reproduce the experimental data for 17°C treatments with admixed nuclear genomic background (we later discuss why this might be), we highlight results for populations with pure Melbourne or pure Townsville nuclear backgrounds; admixed results for 27°C populations are presented in the supplementary material.

The data across all treatments are consistent with A1 having a consistent advantage over B1 when A1 is rare. The posterior mean and credible interval ranges for *V* are consistently greater than one, with A1 having rare-type advantage of roughly 30% to 75%, on average (Fig. 3, with the mean of the posterior for *V* ranges from ∼1.3 to ∼1.75). There is also compelling evidence for negative frequency-dependence of relative fitness, with posterior means and CI for *s* consistently negative, which is consistent with the qualitative observation that A1 often increases a lot when rare and marginally when common (Fig. 2).

**Figure 3:**
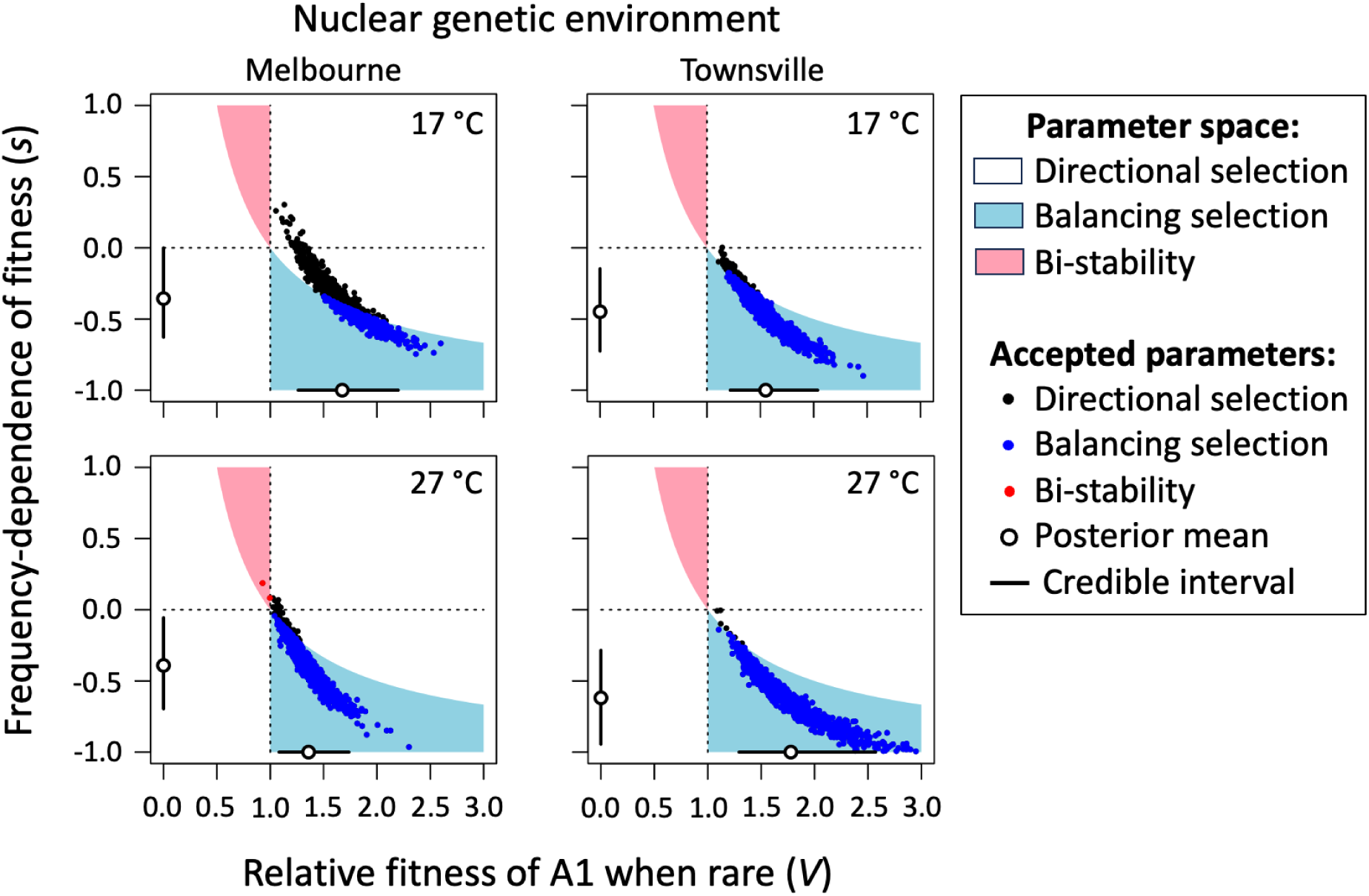
Selection scenarios that are consistent with the evolutionary dynamics of mtDNA haplotypes. Posterior estimates of the ABC analysis are shown for the two selection parameters, *V* and *s*. Each panel shows the entire parameter space over which simulations were carried out (priors for the parameters are independent and uniformly distributed across this space, with -1 < *s* < 1 and 0 < *V* < 3). The broken vertical line delineates the boundary between a rare-A1 haplotype advantage (*V* > 1) and disadvantage (*V* < 1). The broken horizontal line divides positive frequency-dependence (*s* > 0) and negative frequency-dependence (*s* < 0). The filled circles (10^3^ accepted parameter combinations per panel) show simulated values from the posterior parameter distributions (*i.e.*, the “accepted” parameters that were consistent with our actual experimental data, shown in Fig. 2). Open circles and whiskers show the posterior mean and credible interval for each parameter.

Across the entire parameter space our model, there are three basic domains of selection (shown in the light blue, pink, and white shaded regions of Fig. 3): (1) directional selection, in which one of the mtDNA haplotypes exhibits a consistent advantage and selection favours its fixation, (2) balancing selection, which favours the maintenance of both haplotypes, and (3) “bi-stability” in which both fixation states (A1 or B1 fixed) are stable and the long-term fate of each haplotype is highly sensitive to their initial frequencies. The ABC analysis suggests that the experimental data are largely incompatible with the bi-stability scenario. Instead, selection is predominantly directional, with fixation of the A1 haplotype favoured, and balancing (Fig. 3). Directional selection predominates in populations evolving at 17°C with Melbourne nuclear backgrounds, and otherwise selection is balancing (Fig. 4; this includes the 27°C treatments with admixed nuclear backgrounds: see Fig. S3).

**Figure 4:**
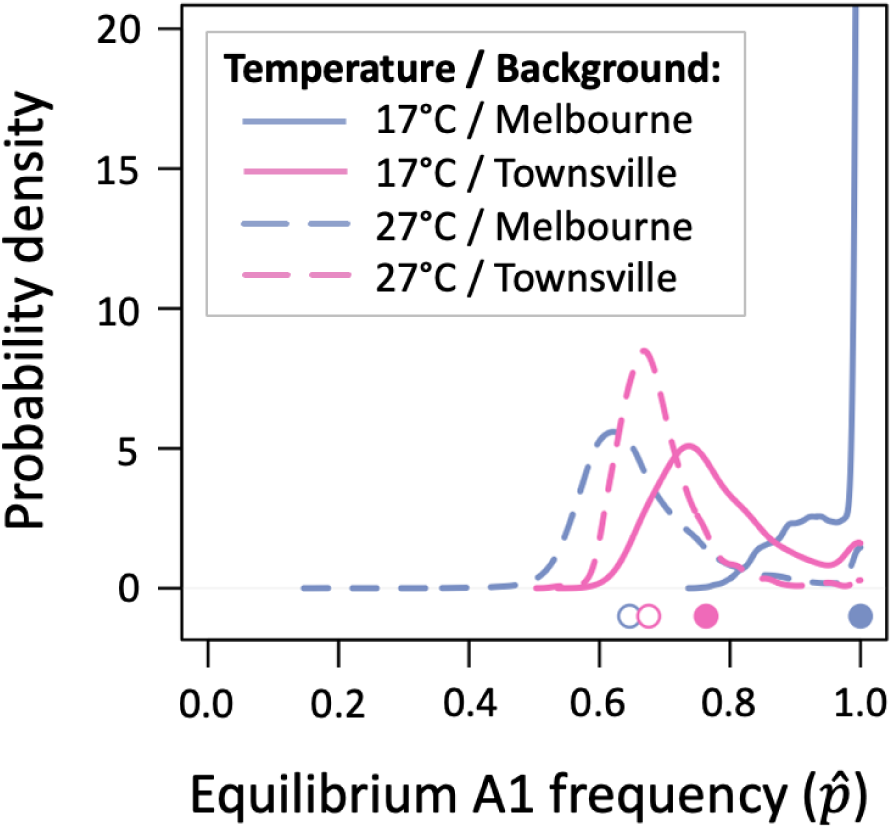
Predicted equilibrium frequencies of the mtDNA haplotypes. The The long-term (equilibrium) mitochondrial haplotype frequencies depend on the parameters of the model (*s* and *V*), with A1 predicted to evolve towards a stable polymorphic equilibrium *𝑝̂* = (1 − 𝑉)⁄𝑠𝑉 under the parameter condition 1 < 𝑉 < 1⁄(1 + 𝑠), and A1 predicted to reach fixation (the single stable equilibrium is *𝑝̂* = 1) under the condition 1 < 𝑉 > 1⁄(1 + 𝑠). The curves show the probability densities for the predicted values of stable equilibria under each thermal condition and non-admixed nuclear genomic background. Values of *𝑝̂* for each treatment were based on 10^4^ accepted values of *s* and *V*, with rare cases of bi-stability excluded from the analysis. The circles represent median values of *𝑝̂* for the four treatments.

## Discussion

Studies over the past three decades have increasingly focused on, and found evidence for, effects of mtDNA genetic variation on fitness components (Rand 2001; Dowling *et al*. 2008; Ballard and Pichaud 2014) Yet, the specific ways in which selection shapes variation and divergence in the mitochondrial genome remain unclear. Recent research has highlighted the potential contributions of both directional selection, driven by environmental factors such as temperature divergence, and negative frequency-dependent selection (NFDS) to the evolution of mitochondrial haplotype frequencies (Dowling and Wolff 2023). In this study, we used an experimental framework designed to disentangle the contributions of each, by controlling the nuclear genomic backgrounds and initial mtDNA haplotype frequencies of each experimental population, and the environmental conditions in which it evolves. Recognising that the relationship between mtDNA haplotype and phenotype could be modulated by the nuclear genetic background, we predicted that evolutionary responses of mitochondrial haplotypes would vary across the nuclear backgrounds of the populations.

Despite the small size of the mitochondrial genome, our results demonstrate consistent and often strong selection for the A1 haplotype. The A1 and B1 haplotypes are well studied and segregate at high frequencies in natural populations of *D. melanogaster* (Camus *et al*. 2017; Lajbner *et al*. 2018). Our results show that these haplotypes strongly affect fitness, and that the advantage of A1 relative to B1 is negative frequency dependent. The A1 haplotype is consistently favoured when rare, and our ABC simulations indicate that negative frequency dependence should lead to balancing selection in most of the thermal environments and nuclear genomic backgrounds we evaluated.

Traditionally, studies exploring the evolutionary trajectories of mitochondrial haplotypes in *Drosophila* have been conducted in population cages, where large numbers of flies with known starting frequencies of mtDNA haplotypes are introduced, and the subsequent frequency changes of the haplotypes monitored over multiple generations. In some experiments, researchers have attempted to replicate observed frequency changes through ‘perturbation’, where haplotype frequencies are intentionally altered and then monitored to see if they return to their pre-perturbation states (MacRae and Anderson 1988; James and Ballard 2003; Mossman *et al*. 2019). In some of these studies, mtDNA haplotypes have been observed to rapidly fix over a relatively short number of generations (Fos *et al*. 1990; Hutter and Rand 1995; García-Martínez *et al*. 1998), while in others haplotype frequencies converged towards relatively stable polymorphic frequencies, consistent with NFDS (MacRae and Anderson 1988; Hutter and Rand 1995). These prior results are notable in the context of our study, as we observed that the A1 haplotype generally increased in frequency, though much more so from a low initial frequency, which again is consistent with NFDS.

Exceptions were observed in experimental populations with admixed nuclear genomic backgrounds, where A1 haplotype frequencies decreased overall within the cooler 17°C environment. The evolutionary dynamics of A1 in these treatments were incompatible with our simple population genetic model (the basis of our ABC results), which ignores the possibility of mito-nuclear epistatic interactions for fitness. Strong mito-nuclear epistasis would presumably require both interacting loci (assuming pairwise epistasis) to be segregating for intermediate frequency alleles, which is perhaps more likely in the admixed experimental populations where heterozygosity at key nuclear genomic loci is artificially inflated relative to heterozygosity levels in nature (e.g., if the Townsville and Melbourne nuclear backgrounds are nearly fixed for different alleles at the key loci). This possibility could be tested in future studies that use a similar design that compares homogenous and admixed genomic backgrounds but also includes sequencing to enable estimation of nuclear genomic allele frequencies. The specific cause of the interaction between mitochondrial haplotype evolution, admixed nuclear genomic backgrounds, and the cool temperature treatment warrants further study.

Our findings align with a growing recognition that mitochondrial responses to selection are highly context-specific, and often sensitive to both environmental conditions and population genetic backgrounds. Temperature has been identified as a key environmental factor affecting directional selection of mtDNA haplotype frequencies (Mishmar *et al*. 2003; Balloux *et al*. 2009; Lamb *et al*. 2018). Earlier studies suggested that A1-carrying flies exhibit superior tolerance to heat stress and warmer environments (Camus *et al*. 2017; Lajbner *et al*. 2018). These findings were further supported by a quantitative genetic study by Lasne *et al*. (2019), which uncovered a mitochondrial genotypic contribution to adaptive divergence in heat stress tolerance between cline-end Australian populations of *D. melanogaster*—a result likely to be driven by thermotolerance differences between the northern-predominant A1 haplotype and southern-predominant haplotypes (Lasne *et al*. 2019).

However, recent studies challenge this relatively simple story of thermal adaptation mediated by the A1 and B1 haplotypes. Bettinazzi *et al*. (2024) found no differences in thermotolerance between the A1 and B1 haplotypes when introgressed into homogeneous mass bred backgrounds from Melbourne or Townsville populations. Similarly, our findings deviate from earlier results, as we documented a consistent evolutionary advantage to A1 over B1 (outlier admixed results notwithstanding) in both thermal regimes in our study, including stronger selection for A1 at the cooler temperature. These results suggest that the thermal environment is not the only environmental variable that matters for the evolution of A1 and B1 haplotype clines along eastern Australia. Although the clines observed in nature clearly imply that A1 is favoured in subtropical environments and B1 is favoured in temperate environments (Camus *et al*. 2017), cold and heat tolerance cannot be the exclusive determinants of fitness in these local environments. Thermal conditions are likely to contribute to overall fitness, yet there are clearly additional environmental variables that should be considered. Other potential environmental variables that correlate with local thermal conditions include desiccation stress, food content and availability, and interactions with other organisms. These possibilities warrant further investigation.

One intriguing possibility is that endosymbionts interact with the mitochondria to affect fitness. Lajbner *et al*. (2018) carried out experimental evolution of the A1 and B1 haplotypes in mass bred populations under divergent thermal selection (Lajbner *et al*. 2018). In contrast to our results, they identified a consistent trend for A1 to outcompete B1 in warmer conditions, and to be outcompeted in cooler conditions. Interestingly, they replicated their experiments under conditions in which populations had first been purged of *Wolbachia* infection via tetracycline treatment, and conditions in which *Wolbachia* was present. They only identified signatures of adaptive mtDNA responses to thermal selection in the absence of *Wolbachia*, suggesting that *Wolbachia*, which co-segregates with mitochondria in nature, may interact with the mtDNA haplotypes, or erode the capacity of selection to target mitochondrial haplotypes. Notwithstanding, their result highlights yet another factor that may shape responses of selection on mitochondrial haplotypes. Our populations were also free from *Wolbachia* infection, raising the question of what factors might explain the different findings between our study and that of Lajbner *et al*. (2018). These discrepancies are likely caused by differences in the designs of the two studies, which were extensive, spanning the temperatures used to the methods of strain creation. For example, logistical choices in our experimental design may have inadvertently selected for the A1 haplotype in cooler conditions. In our experiment, females oviposited for two days at 27°C and four days at 17°C, at each generation, effectively selecting for high female fertility during early adulthood. This may have inadvertently favoured the A1 haplotype if A1-carrying females consistently outproduced B1-carrying flies in early adulthood, with this discrepancy waning with age. That is, tight regulation of oviposition in our study may have inadvertently resulted in selection for A1, possibly overriding selection mediated by the different temperature treatments.

Our study demonstrated strong and consistent selection for the A1 haplotype across varying conditions of temperature. The results were consistent with strong NFDS, which in some of our treatments would likely lead to balancing selection that maintains polymorphism of the two major mtDNA haplotypes. Our results contrast with previous findings and suggest that ecological factors not represented in our study—such as fluctuating temperatures, resource availability, and biotic interactions—additionally influence mitochondrial evolutionary dynamics in nature. We propose that future laboratory studies move towards incorporating these complexities into their designs to better capture the dynamics of mitochondrial adaptation in natural populations.

In conclusion, we carried out a highly replicated experimental evolution study that tracked the frequencies of two well-studied mtDNA haplotypes sourced from locally adapted populations sampled from opposite ends of an Australian latitudinal cline. Under the controlled conditions of these experiments, and supported by population genetic simulations, we show that A1 is consistently and strongly favoured over B1, particularly when rare, and that this suggests its advantage is negative frequency dependent leading to balancing selection on the haplotype frequencies in most of the nuclear genomic background and temperature contexts studied. Thus, these naturally segregating mtDNA haplotypes have strong, albeit context-dependent, fitness effects, inconsistent with nonadaptive (neutral) explanations for clinal divergence and the maintenance of genetic variation. Nevertheless, our controlled and replicated experimental conditions clearly do not approximate the full spectrum of environmental conditions that distinguish cline-end populations of *D. melanogaster*, and we encourage that future studies consider the additional environmental variables and fitness components (e.g., selection via viability versus fertility) that determine mtDNA haplotypes frequencies in the wild.

## Acknowledgements

We thank Dr. Winston Yee for his help in creating the strains, and help with the initial experimental set-up and in maintaining populations throughout the experimental evolution. We also thank Dr. Travis K Johnson for his help with primer design for differentiating between the A1/B1 haplotypes. This work was supported by funding from the Australian Research Council (DP210102931).

## Author Contributions

DKD acquired the project funding; JRK, DKD and TC designed the study; JRK and ANA performed the experimental evolution and collected the data; JRK, DKD and ANA conducted the statistical analyses of the experimental evolution data; TC designed and conducted the population genetic modelling; JRK, DKD and TC interpreted and synthesised results, JRK wrote the first draft of the manuscript and JRK, DKD and TC iterated subsequent drafts; all authors read and provided input on the final draft.

