## Supplementary information for "Dissecting contributions of directional and balancing selection to trajectories of mitochondrial haplotype evolution in *Drosophila melanogaster*"

**Table S1: Combinations and proportions of mtDNA haplotypes and population backgrounds within the experimental populations subject to experimental evolution.** Experimental populations were seeded with different starting frequencies of the A1 haplotype (either 30% or 70%). Each population was also comprised one of four population backgrounds; either a homogenous Townsville (denoted in pink) or Melbourne (green) nuclear background, or one of two heterogeneous nuclear backgrounds which consist of different proportions of the Townsville nuclear background; either 30% Townsville (orange) or 70% Townsville (purple) background.

| | **Haplotype and Nuclear Background contributing 70% of the population** | | | | |
|---|---|---|---|---|---|
| **Haplotype and Nuclear Background contributing 30% of the population** | | A1 Towns | A1 Mel | B1 Towns | B1 Mel |
| | A1 Towns | | | 30% A1 Towns x 70% B1 Towns | 30% A1 Towns x 70% B1 Mel |
| | A1 Mel | | | 30% A1 Mel x 70% B1 Towns | 30% A1 Mel x 70% B1 Mel |
| | B1 Towns | 70% A1 Towns x 30% B1 Towns | 70% A1 Mel x 30% B1 Towns | | |

| | | | |
|---|---|---|---|
| | B1 Mel | 70% A1 Towns x 30% B1 Mel | 70% A1 Mel x 30% B1 Mel |

Cross 1: ♀ Wildtype mitochondrial strain (mt) x ♂ FM7/Y → ♀ FM7/+(mt)

Cross 2: ♀ FM7/+(mt) x ♂ Target Nuclear Background (nuc) → ♀ nuc/FM7(mt)

Cross 3a: ♀ nuc/FM7(mt) x ♂ nuc → ♀ nuc/nuc; +/+; +/+(mt)

Cross 3b: ♀ nuc x ♂ CyO/+; TM6B/+ → ♂ nuc/Y; CyO/nuc; TM6B/nuc

Cross 4: ♀ nuc/nuc; +/+; +/+(mt) x ♂ nuc/Y; CyO/nuc; TM6B/nuc → ♀ nuc/nuc; CyO/+; TM6B/+(mt)

Cross 5: ♀ nuc/nuc; CyO/+; TM6B/+(mt) x ♂ nuc/Y; nuc/nuc; nuc/nuc → ♀ nuc/nuc; CyO/nuc; TM6B/nuc(mt)

Cross 6: ♀ nuc/nuc; CyO/nuc; TM6B/nuc(mt) x ♂ nuc/nucY; nuc/nuc; nuc/nuc → ♀ + ♂ nuc X and Y; nuc/nuc; nuc;nuc(mt)

**Figure S1:** Crossing scheme used to introgress the Melbourne/Townsville nuclear backgrounds into the A1 and B1 haplotype strains. mt = mitochondrial haplotype (either A1 or B1), nuc = target nuclear background of a given chromosome where FM7 is an X chromosome balancer, CyO is a second chromosome balancer and TM6B is a third chromosome balancer. Target genotypes resulting from each cross are denoted on the right side of each cross. This method is based on the scheme described in Clancy (2008).

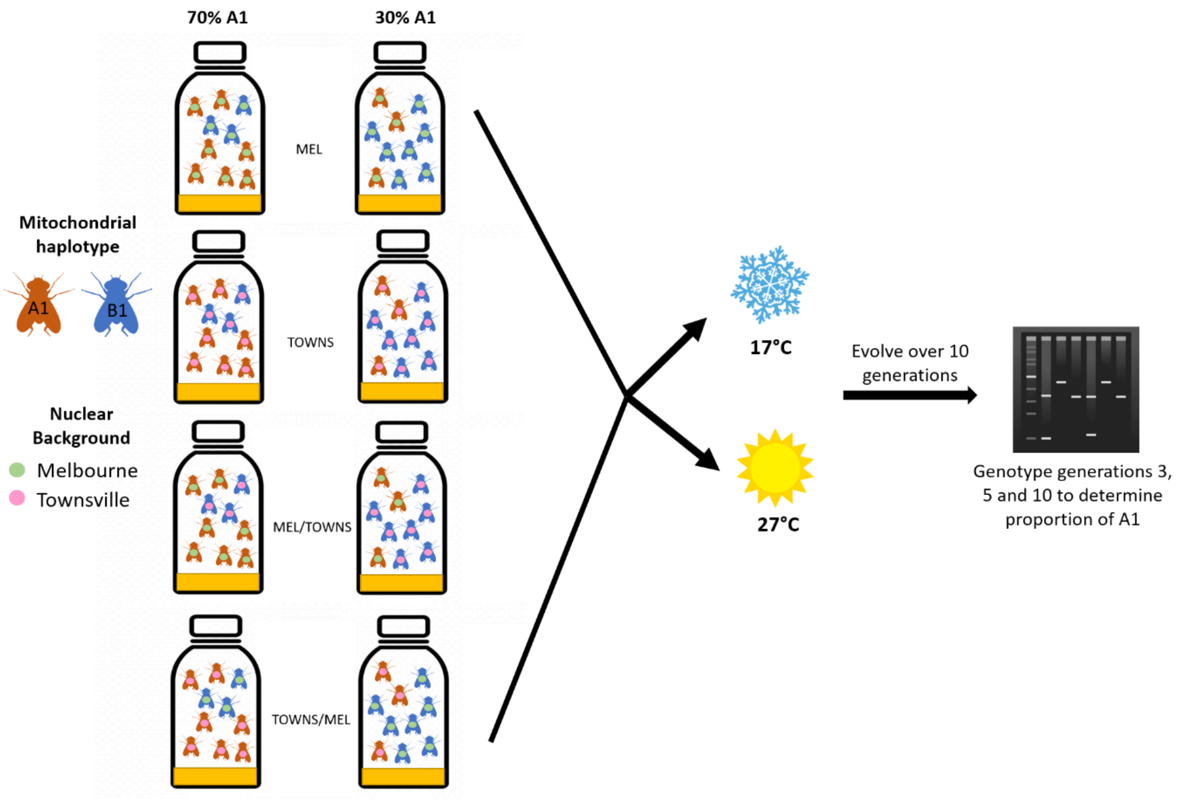

**Figure S2: Experimental evolution design.** Populations with either an A1 starting frequency of 30% or 70% were set up so that the populations comprised one of four population backgrounds; either a homogenous Melbourne or Townsville background, or either a heterogeneous 30% Townsville or 70% Townsville nuclear background. Populations were then evolved at either 17°C or 27°C for 10 generations after which the frequency of the A1 haplotype was determined by genotyping 30 female flies from each population at generations 3, 5 and 10.

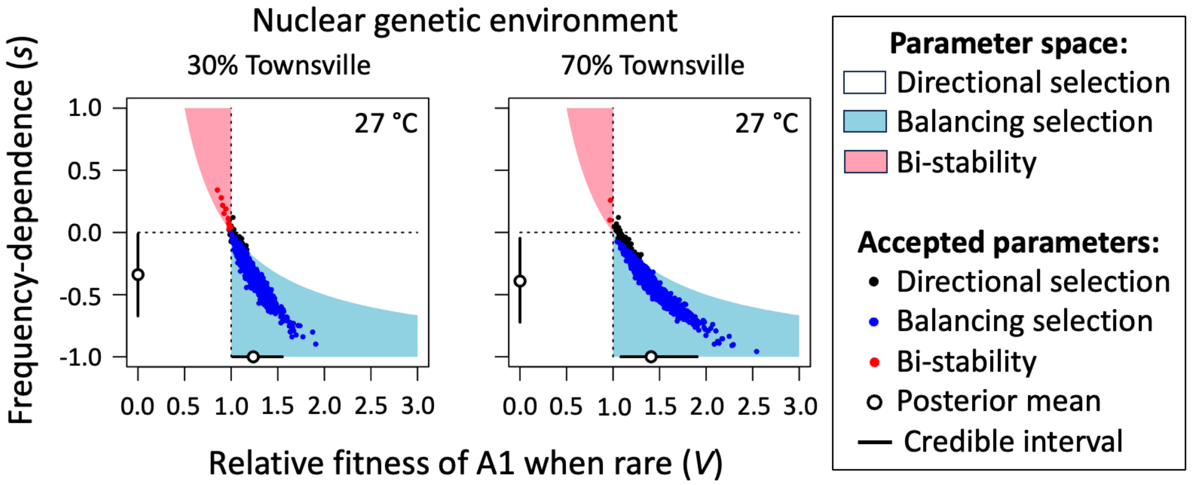

**Figure S3: ABC analysis of populations with admixed nuclear backgrounds in 27°C environments.** Posterior estimates of the ABC analysis are shown for the two selection parameters, $V$ and $s$. Each panel shows the entire parameter space over which simulations were carried out (priors for the parameters are independent and uniformly distributed across this space, with $-1 < s < 1$ and $0 < V < 3$). The broken vertical line delineates the boundary between a rare-A1 haplotype advantage ($V > 1$) and disadvantage ($V < 1$). The broken horizontal line divides positive frequency-dependence ($s > 0$) and negative frequency-dependence ($s < 0$). The filled circles show simulated values from the posterior parameter distributions (*i.e.*, the "accepted" parameters that were consistent with our actual experimental data, shown in Fig. 2). Open circles and whiskers show the posterior mean and credible interval for each parameter.
